# Hidden Markov Models Lead to Higher Resolution Maps of Mutation Signature Activity in Cancer

**DOI:** 10.1101/392639

**Authors:** Xiaoqing Huang, Itay Sason, Damian Wojtowicz, Yoo-Ah Kim, Mark D.M. Leiserson, Teresa M. Przytycka, Roded Sharan

**Author notes:** Equal first author contribution. Equal senior author contribution (Itay Sason, Damian Wojtowicz, Mark D.M. Leiserson, Teresa M. Przytycka, Roded Sharan).

## Abstract

Knowing the activity of the mutational processes shaping a cancer genome may provide insight into tumorigenesis and personalized therapy. It is thus important to uncover the characteristic *signatures* of active mutational processes in patients from their patterns of single base substitutions. However, mutational processes do not act uniformly on the genome and are biased by factors such as the genome’s chromatin structure or replication origins. These factors may lead to statistical dependencies among neighboring mutations, calling for modeling approaches that can account for such dependencies to better estimate mutational process activities.

Here we develop the first sequence-dependent models for mutation signatures. We apply these models to characterize genomic and other factors that influence the activity of previously validated mutation signatures in breast cancer. We find that our tool, SigMa, can accurately assign genomic mutations to mutation signatures, yielding assignments that are of higher likelihood than those obtained with models that assume independence between signatures and align better with current biological knowledge. Our analysis resolves a controversy related to the dependency of APOBEC signatures on replication time and links Signatures 18 and 30 to oxidative damage.

Modeling the sequential dependencies of mutation signatures leads to improved estimates of mutation signature activity both at the tumor-level and within specific genomic regions, yielding higher resolution maps of mutation signature activity in cancer.

## 1 Introduction

Cells acquire somatic mutations over time from exposure to different combinations of mutational processes, potentially leading to cancer. Understanding the activity of mutational processes is critical for cancer treatment, as many standard treatments introduce DNA damage or inhibit DNA damage repair genes [8, 19]. Presently, clinicians use specialized assays for specific biomarkers to characterize DNA damage repair deficiencies, such as microsatellite instability (see, e.g., [11]). Large-scale cancer sequencing efforts have recently opened up new avenues for characterizing the activity of mutational processes. The key insight is that mutational processes leave *signatures* of their activity in cancer genomes, the most well-studied of which are patterns of base substitutions.

An increasing body of research aims at inferring signatures and their exposures from large datasets of mutations from cancer whole-exome and -genome sequences [3, 4, 17, 28, 44, 42, 25], and the Catalogue of Somatic Mutations in Cancer (COSMIC) consortium has collected a census of 30 validated mutation signatures [18]. Many of these signatures are associated with deficient DNA damage repair pathways; some have been validated experimentally [15, 48], expanding the opportunity for targeted therapy. For example, Davies et al. [12] provided evidence that mutation signatures reveal patients deficient in homologous recombination repair (HR), and thus may benefit from PARP inhibitor treatment. Importantly, some of these patients do not harbor biallelic inactivations in known HR genes. Other signatures are associated with environmental exposures to carcinogens such as tobacco smoke [2] or aflatoxin [35], and two are associated with aging [1] indicating that the underlying mutational processes may be active in healthy cells.

Despite these advances, uncovering etiology of mutation signatures and inferring their exposures remain significant challenges, e.g. about half of the COSMIC signatures have no known etiology. Even with validated mutation signatures, it can be difficult to infer their exposures and assign individual mutations to the corresponding signature, in part because there may be multiple signatures of the same mutational process. One key factor to inferring signature exposure is the sequential dependency of the signatures. This is the idea that mutations that are adjacent in a given cancer genome are more likely to be the result of the same mutation signature. In their seminal work, Nik-Zainal et al. [33] identified clusters of mutations in breast cancers (termed *kataegis*) that display a particular base substitution signature. Kasar et al. [28] uncovered a signature of “canonical” activation-induced cytidine deaminase (AID) pathway activity in chronic lymphocytic leukemia that was missed by Alexandrov, et al. [3]. Part of the reason for their discovery was that they incorporated the “nearest mutation distance” into their model, since AID is known to cause multiple mutations within local regions of the genome. Morganella et al. [31] identified so-called “processive groups” of up to 20 mutations believed to come from the same signature. Morganella et al. [31] and Haradhvala et al. [22] both characterized signatures in terms of the transcriptional and replicative strands, and replication timing. Supek and Lehner [45] identified mutation signatures that are specifically associated with clusters of mutations, and showed that the activity of these signatures is associated with an increase in the mutation rate of expressed genes.

Motivated by this earlier work, we set out to model the genomic factors that bias mutational process activity, such as genome position, CpG islands, and replication origins. We hypothesized that by capturing the statistical dependencies introduced by these genomic factors, our models would yield more precise estimates of mutation signature exposure, and would further reveal genomic features that correlate with mutational process activities. Our contribution is three-fold: (i) we suggest the first probabilistic model to account for sequential dependency among mutation signatures; (ii) we use this model to rigorously assign mutation signatures to individual mutations and characterize the genomic and phenotypic preferences of mutation signatures; and (iii) we study the transition probabilities between different mutation signatures.

## 2 Methods

### 2.1 A Hidden Markov Model of mutation signatures

Following previous work, we categorize mutations in a cancer genome into *L* = 96 categories that include its base substitution (C:G>A:T, C:G>T:A, C:G>G:C, A:T>C:G, A:T>T:A, A:T>G:C), and left-(4) and right-flanking (4) nucleotides [3]. We model an observed sequence of mutations using a multinomial Hidden Markov Model (HMM). The model assumes that each observation, representing a mutation category, is emitted by one of *K* states in a Markov chain, representing a mutation signature. The sequence of states that generated the observed sequence is unknown, but as the states form a Markov chain, each state depends on the previous state, thus capturing sequential dependencies between states. An HMM is parameterized by a vector π of *K* starting probabilities, a *K × K* transition matrix *A*, and a *K × L* emission matrix *E*.

The above HMM can capture sequential dependencies but is less motivated for “isolated” mutations that are distant from any other mutation. We call such mutation regions *sky* and refer to regions of close-by mutations as *clouds* (using a distance threshold of 2000 bp as explained below).

We model sky mutations using a multinomial mixture model (MMM). The MMM is characterized by a vector *g* of *K* mutation signature marginal probabilities and the same emission matrix *E*. To model cloud mutations, we use a dynamic Bayesian network (DBN) that is a simple extension of an HMM in that it allows subsequences generated by the HMM to be interspersed with mutations generated by the MMM (for a review of DBNs, see [32]). We call the resulting composite model SigMa (Signature Markov model); a simplified overview of the model is presented in Figure 1 and its cloud component is sketched in Figure S1.

**Fig. 1.**
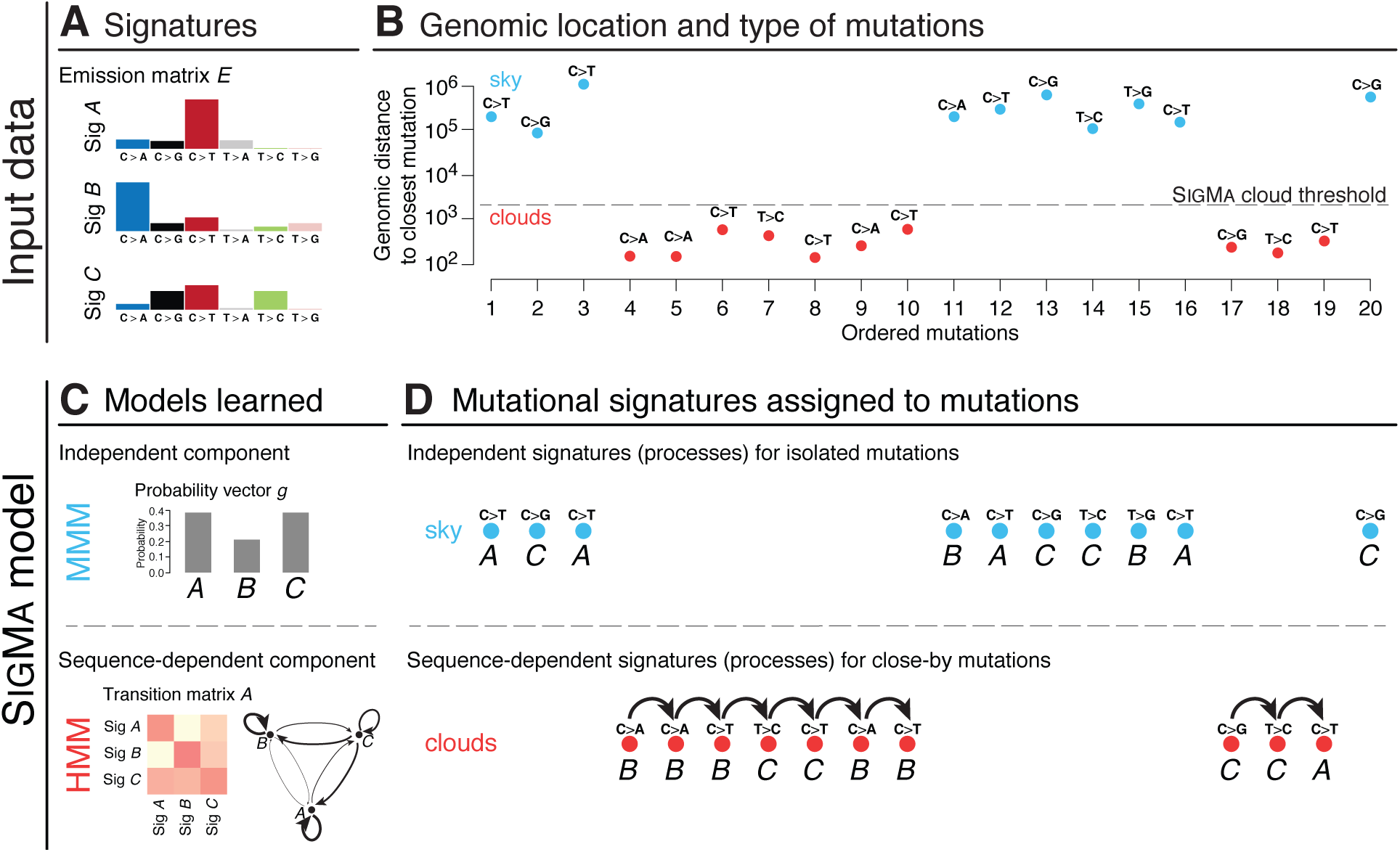
Overview of the SigMa model. The input data consists of (**A**) a set of predefined signatures that form an emission matrix *E* (here, for simplicity, represented over six mutation types), and (**B**) a sequence of mutation categories from a single sample and a distance threshold separating sky and cloud mutation segments. (**C**) The SigMa model has two components: (top) a multinomial mixture model (MMM) for isolated sky mutations and (bottom) an extension of a Hidden Markov Model (HMM) capturing sequential dependencies between close-by cloud mutations; all model parameters are learned from the input data in an unsupervised manner. (**D**) SigMa finds the most likely sequence of signatures that explains the observed mutations in sky and clouds.

We now define the SigMa model for clouds. The input data is a sequence of *O*_1_*,…, O_T_* mutation categories. The (hidden) signature that generated mutation category *O_t_* is represented by *Q_t_*. The transitions between signatures at each subsequent position depend on whether the observed mutation category occurs within sky (marked by a binary indicator *I_t_*) or clouds. The joint probability distribution of the model is:

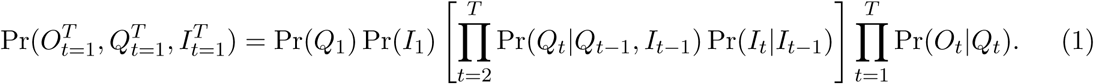

We now define the conditional probability distributions (CPDs). The transition between signature states *Q_t−_*_1_ to *Q_t_* depends on the indicator *I_t−_*_1_ in the following manner. Within sky, the transitions occur according to the marginal probability of each state (i.e., as in the MMM), while otherwise the transitions to state *Q_t_* depend on state *Q_t-_*_1_. Formally, when *I_t_* = 0 (i.e., the current mutation is in a cloud):

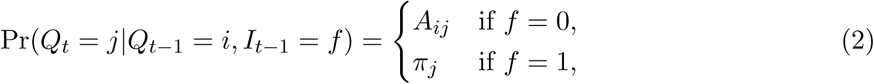

The probability of the initial state depends only on the starting probabilities of the signatures, such that Pr(*Q*_1_ = *i*) = *π_i_*.

The transitions between the sky segment indicator *I_t_* only depends on the previous indicator *I_t-_*_1_, i.e.

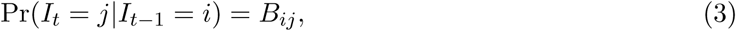

where *B* is the 2 *×* 2 transition matrix between sky and cloud segments. Note that *B* implicitly governs the length of those segments. The probability of starting in a sky/cloud state is given by Pr(*I*_1_ = *i*) = *ρ_i_*, where *ρ* is a 2 *×* 1 starting probability vector. Finally, given the state *Q_t_*, each observation *O_t_* is independent of all other variables, i.e.

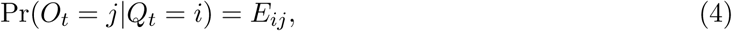

### 2.2 Model training

We learn the SigMa model parameters from data using the Baum-Welch expectation-maximization algorithm with random initialization. We then compute Viterbi paths – the most likely sequence of states that generated the data – to assign mutations to signatures and compute signature *exposures* (i.e. signature frequency per sample). In practice, we find that the assignments are robust with respect to the random initialization used in the learning process; over 99.5% of mutations are assigned to the same signature when compared to majority assignments in 100 random initialization runs of SigMa.

Rather than model the mutations in a cohort of cancer genomes with a single SigMa, we train a model per sample. The motivation for this approach comes from the assumptions of earlier methods (e.g. [4]) that signature exposures are different across samples.

The SigMa model has several meta-parameters that are set in advance: (i) the set of signatures used; and (ii) a distance threshold indicating the beginning of a new segment (cloud or sky) of mutations. In this work we focus on assignment of signatures to mutations rather than on signature learning; hence, we consider only COSMIC signatures ^4^, focusing on the signatures previously identified as active in breast cancers: Signatures 1, 2, 3, 5, 6, 8, 13, 17, 18, 20, 26, and 30. For the other meta parameter, we perform model selection and evaluate the performance of each choice using the log-likelihood of the model on held-out data. To this end, we use a leave-one-out cross-validation scheme leaving one of the chromosomes out. The results are summarized in Figure 2 and accordingly we set the distance threshold to 2000 bp at which the held-out log-likelihood was maximized. Thus, a mutation whose flanking mutations (if any) are more than 2000 bp away is called *sky*; otherwise, the mutation is considered to be within a *cloud*.

**Fig. 2.**
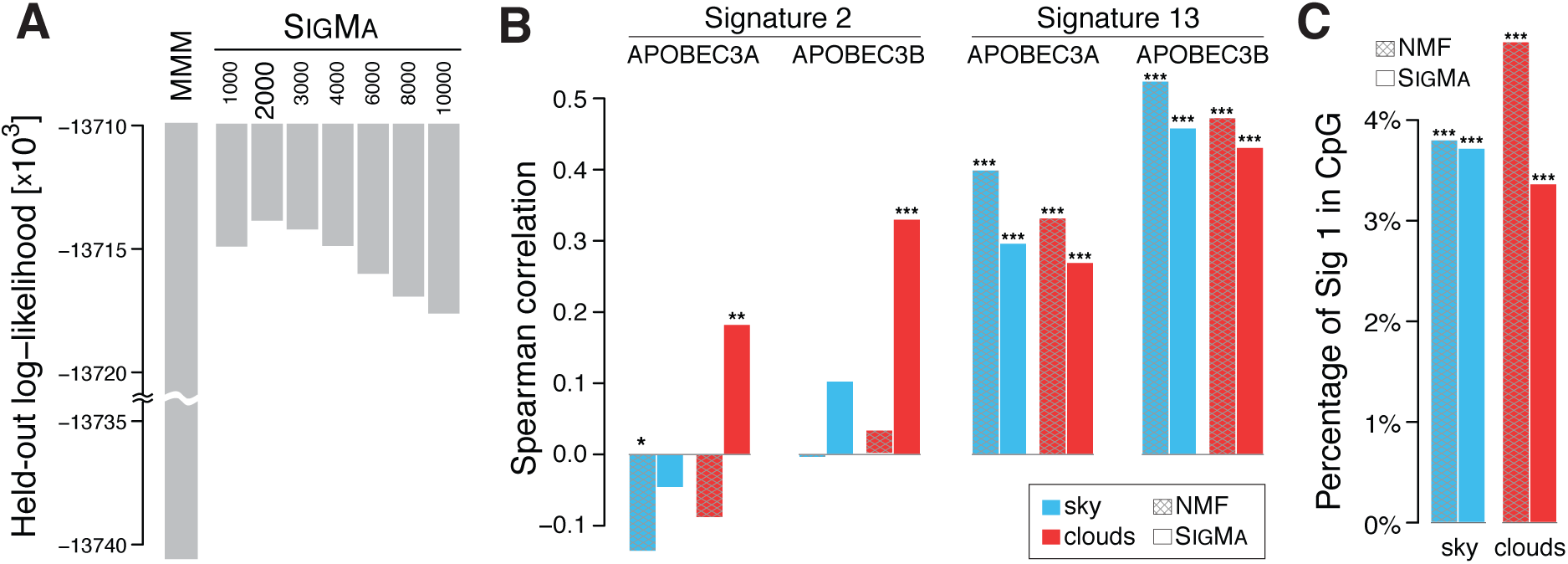
(**A**) Comparative assessment of model performance on held-out data for MMM and SigMa across different distance thresholds. SigMa at a threshold of 2000 bp shows the best performance by maximizing the log-likelihood (the y-axis has a customized scale with a scale break). (**B**) Spearman correlation comparison of APOBEC3A/B expression with Signature 2 and 13 activities across samples. For Signature 2, the mutation counts in clouds with SigMa are positively correlated with APOBEC3A/B expression while the NMF-based counts have zero or negative correlation in both sky and clouds. Signature 13 mutation counts are positively correlated in both models. (**C**) Comparison of fraction of Signature 1 mutations found in CpG islands in sky and clouds. Both NMF and SigMa show significant depletion of Signature 1 in CpG islands with respect to randomized data, with SigMa exhibiting more pronounced depletions, particularly in clouds. We performed 1000 permutations of signature assignments preserving mutation trinucleotide context within each sample. We used a one-sided Wilcoxon signed rank test to compare the observed and randomized numbers of Signature 1 in CpG islands. In (B) and (C), the significance level was categorized as: * P-value (*P*) *<* 0.05; ** *P <* 5 *×* 10^−3^; *** *P <* 5 *×* 10^*−*5^.

We note that while SigMa always models mutations in sky as being independent from one another, the Markovian component of SigMa learns whether the mutations in clouds are sequence dependent or independent. In practice, approximately 88% of mutations in clouds are found to be sequence dependent according to the most likely sequence of mutation events.

### 2.3 Software availability

We implemented SigMa in Python 3. The code is publicly available at https://github.com/lrgr/SIGMA. On average, it takes approximately 8 seconds to train SIGMA on a breast cancer whole-genome, learning a total of 146 model parameters.

### 2.4 Data

We analyzed 3,479,652 mutations in the cohort of 560 breast cancer (BRCA) whole-genomes previously analyzed by Nik-Zainal et al. [34]. Each patient has an average of 208.2 clouds containing an average of 2.33 mutations, with 271,492 total mutations in clouds (8%) and 3,208,160 total mutations in sky (92%).

To compare SigMa to NMF, we recomputed the NMF assignments of signatures to mutations used by Morganella et al. [31] following their maximum likelihood approach, and used these to compute exposures. We downloaded the gene expression data for 266 BRCA samples from Table S7 in Nik-Zainal et al. [34]. For replication timing analysis, we downloaded percentage normalized replication time estimates from Repli-seq data in the MCF-7 cell line from the ENCODE project ^5^ and we split them them into deciles; the MCF-7 cell line was chosen as it most closely represents breast cancers (see Morganella et al. [31] for details). The CpG islands’ coordinates were downloaded from the UCSC Genome Browser.

We evaluate inferred mutation signature exposures in part using clinical and demographic features of each of the 560 cancers. We downloaded clinical and demographic data from Table S1 in Nik-Zainal, et al. [34], restricting our analysis to those features that are measured in at least 85% of the patients (omitting gender since the cohort is *>* 99% female): age, tumor grade, estrogen-receptor (ER) status, progesterone-receptor (PR) status, and HER2 status. We imputed missing data using the mean.

## 3 Results

In order to capture sequential dependencies among mutation signatures we propose a hidden Markov modeling framework. Within this framework, the identity of the mutation signature underlying a given mutation depends (through conditional probability) on the identity of the signature that yielded the preceding mutation in the genome. This modeling approach is motivated by earlier work that has shown that the mutations in localized clusters are often found to be from the same signatures [31, 45], thus suggesting a sequential dependency among mutation signatures. However, the majority of mutations in the cancer genome are hundreds of thousands of base pairs from the nearest mutation, suggesting that this dependency only manifests on small, localized regions of the cancer genome. To account for this complexity, we develop a composite model, SigMa, that can infer the sequential dependencies among mutation signatures within localized densely-mutated regions. We apply SigMa to 560 breast cancer whole-genomes previously analyzed in [34], partitioning each tumor’s mutations into *sky* (isolated mutations) and *clouds* (groups of close-by mutations). The model is sketched in Figure 1; full details on the model, mutation partitioning, and data appear in the Methods.

### 3.1 SigMa uncovers sequential dependency between mutation signatures and leads to stronger associations with related biological signals

To assess the utility of SigMa in capturing sequential dependencies, we compare it to a baseline probabilistic model with no sequential dependencies. Since the state-of-the-art method for inferring mutation signatures, non-negative matrix factorization (NMF), is non-probabilistic, we use a related multinomial mixture model (MMM) as our baseline. The model parameters are learned so as to maximize the likelihood of the model using Expectation Maximization. Both SigMa and MMM were applied to each sample separately, fixing the twelve COSMIC signatures previously found to be active in breast cancer (see Methods for details).

Figure 2A summarizes the performance of the models in cross validation on a breast cancer dataset of 560 genomes. We draw two conclusions from these results. First, there is significant sequential dependency among the mutation signatures within clouds, as the variants of SigMa all outperform the baseline MMM. Second, the sequential dependency is strongest for mutations within 1000-4000 bases of one another, as SigMa achieves the highest held-out log-likelihood in this range using a distance threshold of 2000 bp. Thus, we adopt the threshold of 2000 bp for the remainder of our experiments.

An important feature of our proposed probabilistic model is that it allows inferring the most likely mutation events that led to the observed data. Hence, we wished to assess if the inferred assignments of mutations to signatures can strengthen the associations with related biological signals in comparison to NMF-based assignments [31]. One of the best understood signature is the age-related Signature 1, the result of an endogenous mutational process initiated by spontaneous deamination of 5-methylcytosine. This process occurs at cytosine-guanine (CpG) dinucleotides and is related to the major site of cytosine methylation which carries the risk of spontaneous deamination of 5-methylcytosine (5mC) to yield thymine. CpG methylation is a silencing mark. CpG islands are GC-rich genomic regions that are often located around gene promoters of active genes and are typically not methylated [37, 13]. Thus, we expect a depletion of Signature 1 in those regions after correction for trinucleotide context of mutations. While both models shows significant depletion of Signature 1 in CpG islands, SigMa exhibits more pronounced depletions, especially in clouds (Figure 2C).

APOBEC enzymes are another relatively well understood source of mutations in cancer. The APOBECs deaminate cytosines in single-stranded DNA, preferentially at TpC sequence context and are thus believed to be associated with Signature 2 and 13. In particular, APOBEC3A and APOBEC3B are among the main factors causing mutations in human cancers and specifically implicated in inducing clustered mutations (kataegis) [5, 6, 46], prompting us to test for an association between APOBEC3A/B expression and the number of mutations assigned to Signatures 2 and 13. Surprisingly, the NMF-based mutation assignments show no or negative correlation between Signature 2 and APOBEC3A/B expression both in sky and clouds, and a statistically significant correlation was observed only with Signature 13. In contrast, using mutation assignments from SigMa, we find that the Signature 2 mutation counts in clouds show positive correlations with the APOBEC3A and APOBEC3B expression (*P* = 4.6 *×* 10^*−*3^ and 6.3 *×* 10^*−*8^, respectively; Figure 2B). Signature 13 mutations remain positively correlated in both sky and clouds.

### 3.2 Sky and clouds show distinct mutations patterns

In SigMa, clouds are defined as dense groups of mutations, but unlike the definition of clustered mutations [45] or processive groups [31], we make no restriction on consecutive mutations being of the same type and/or being on the same strand. We also do not require that the number of mutations in a cloud is large or filter out nearby mutations. Despite our liberal 2000 bp cut-off for maximal distance of two constitutive mutations in a cloud, median distances between mutations in the same cloud are less than 500 bp independently of its size (number of mutations in a cloud; see Figure 3A) while the median distance between mutations in the sky is more than 150,000 bp. As expected, the differences in mutation assignments between SigMa and NMF are much higher for the mutations that belong to clouds than to sky (Figure 3B).

**Fig. 3.**
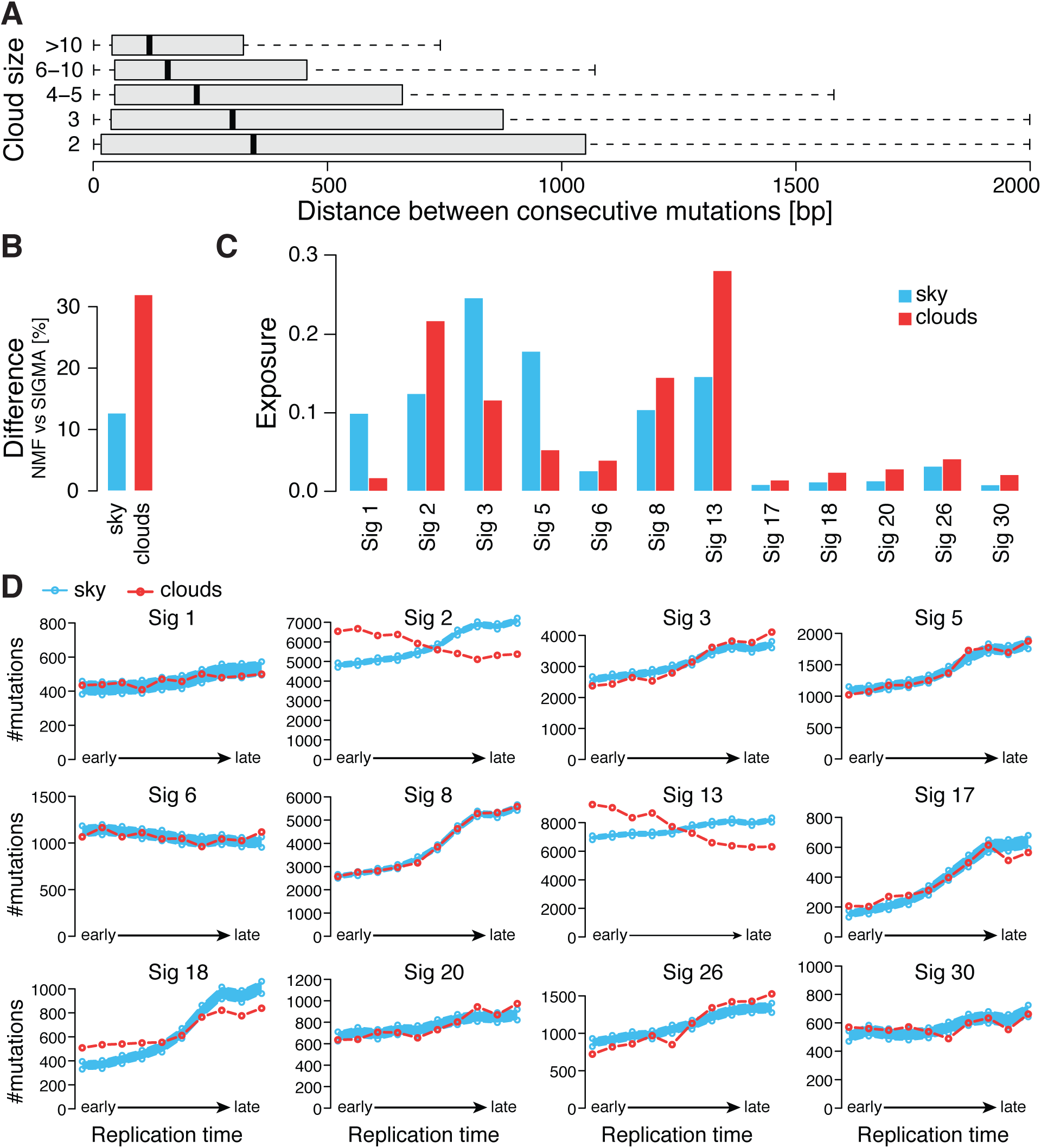
(**A**) Distribution of distance between consecutive mutations in clouds for clouds of various sizes (number of mutations in a cloud). (**B**) Difference between NMF and SigMa in mutation signatures assigned to mutations is higher for cloud mutations. (**C**) Comparison of exposure to mutation signatures in sky and cloud regions based on SigMa signature assignments. (**D**) Frequency distribution of the 12 mutation signatures (assigned by SigMa) over replication time. The red line is the distribution of frequencies over replication time from early to late for mutations in clouds. In blue is a distribution of trends for sky mutations downsampled to the number of mutations found in clouds. The sampling was repeated 1000 times, and the 95 percent confidence intervals of the downsampled sky mutation frequencies over replication time are shown.

Interestingly, clouds and sky show quite different distribution of signature exposures. For example, clouds are strongly enriched in Signatures 2, 8, 13, 30 but depleted in Signatures 1, 3, and 5 (Figure 3C).

The above observations suggest that the properties of clouds and sky are quite different. Moreover, we observed that sky mutations show an about 50% increase of mutations in late replication regions, compared to about a 15% increase in clouds (Figure S2). Therefore, we analyzed the distribution of mutations assigned to individual signatures with respect to replication time considering clouds and sky as two potentially different subpopulations. With the exception of mismatch repair Signature 6, all signatures within sky are enriched in late replication regions (Figure 3D). Some signatures, such as Signatures 1, 5, and 8, show no appreciable differences in the trends between sky and clouds; however, most of the signatures do. The most striking difference in the trends is displayed by the APOBEC Signatures 2 and 13. Previous studies that analyzed the relation of APOBEC with replication time appeared to be contradictory. Kazaonov et al. [29] reported enrichment of APOBEC mutations in early replicating regions and hypothesized that this unusual mutagenesis profile may be associated with a higher propensity to form single-strand DNA substrates for APOBEC enzymes in early-replicating regions. However, Morganella et al. [31] found that Signature 2 is enriched in late replicating regions suggesting that APOBEC mutations assigned to Signature 2 are more efficiently repaired in early replicating regions. They were also surprised to find that Signature 13 differed from Signature 2 and showed no dependency of mutation frequency on replication time (see also Figure S3). Our analysis reconciles these two results and demonstrates that while APOBEC mutations associated with clouds show properties consistent with these reported by Kazaonov et al., the sky associated ones show the usual enrichment in late replicating regions. The cumulative mutation profile depends on the individual characteristics of the sky-associated and clouds-associated subpopulations and their relative abundance. Interestingly, the proportion of cloud-associated mutations relative to sky-associated mutations is higher for Signature 13 than for Signature 2 (Figure 3C) contributing to the differences in cumulative trends of these two signatures reported by Morganella et al. (Figure S3).

Overall, this analysis demonstrates that some signatures have very different properties when considered in the context of clouds versus sky suggesting that the interplay of mutational processes that underline the same signature in sky and in clouds might be different.

### 3.3 Transition probabilities reveal an association between Signatures 30 and 18, linking them to oxidative damage

Next, we asked if the transition probabilities can provide additional insights into etiology of mutation signatures. Since the number of cloud mutations in individual patients is small, we used cumulative transition probabilities obtained by counting the transitions between signatures in clouds across all samples. The most frequent transitions are from each signature to itself (Figure 4A). Nevertheless, there are also frequently occurring transitions between pairs of different signatures; we quantified the enrichment of transition probabilities between signatures using Pearson residuals (Figure 4B).

**Fig. 4.**
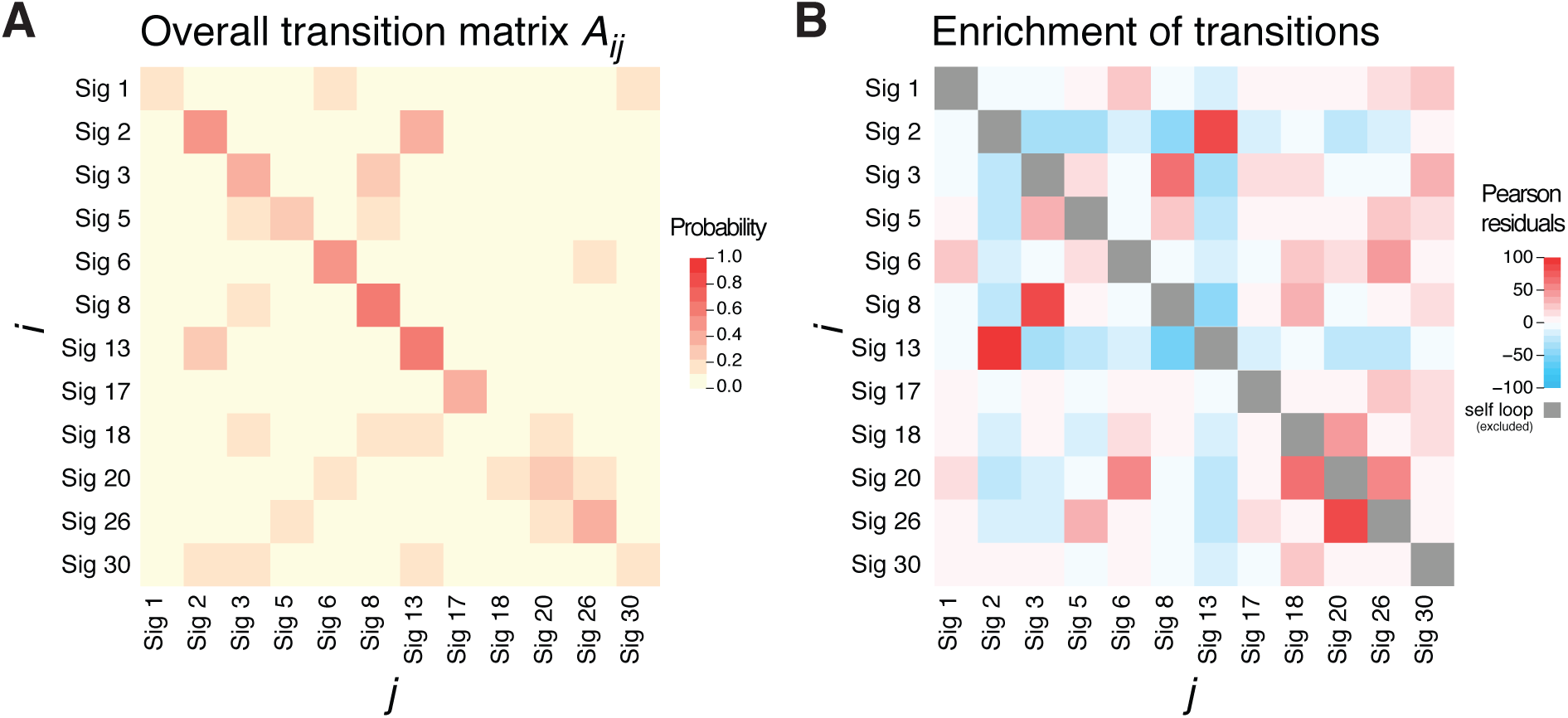
(**A**) Learned transition probabilities between mutation signatures in sequence-dependent cloud segments across all samples. For most signatures, self transitions have high probabilities. **B**) Enrichment of transition frequencies between different signatures represented by Pearson residuals between observed and expected (based on total exposures) frequencies. Since self transition are all enriched (Pearson residuals ranging from 24 (Sig 18) to 168 (Sig 6)) they were excluded from this analysis to correctly estimate enrichment of transitions between different signatures.

Expectedly, we observed an enrichment of transitions between the two APOBEC Signatures 2 and 13 and between the mismatch repair Signatures 6, 20, and 26. These are groups of different (and dissimilar) signatures that are known to be underlined by the same general mutagenic processes and are often found in the same samples. Interestingly, there is also strong association between Signatures 3 and 8 suggesting a relation between Signature 8 and homologous recombination deficiency that was shown to underlie Signature 3 [39].

Surprisingly, we also observed an enrichment in transitions between signatures 18 and 30 suggesting a possible relation between these less understood signatures. Further supporting this relationship, we found that these signatures significantly co-occur in the same patients (*P <* 2.2*×*10^*−*16^ for clouds based on Fisher exact test where signatures with exposure at least 0.01 are considered to be present; co-occurrence is not significant in sky). Previous studies linked a new signature that is very similar to Signature 18 to bialleic deactivation of *MUTYH*, which is involved in base excision repair in response to oxidative damage [47, 43, 36]. Specifically, *MUTYH* is involved in repairing the damage caused by 8-Oxoguanine (8-oxoG) – one of the most common DNA lesions resulting from presence of Reactive Oxygen Species (ROS). If not corrected, it leads to G-to-T transversion.

As for Signature 30, recent studies linked it to mutations in the *NTHL1* gene [15]. Similarly to the *MUTYH* gene, *NTHL1* is a glycosylase that is also involved in the repair of oxidative DNA damage. Unlike *MUTYH* which is involved in the repair of oxidized purines, *NTHL1* is involved in the removal of oxidative pyrimidine lesions. If not corrected, oxidized, deaminated cytosines are a source of C-to-T transitions in vivo [30] which is consistent with the mutational profile of Signature 30.

However, *MUTYH* appears to be functional in breast cancer and while it is natural to assume that Signature 18, which has been observed in breast cancer, is somehow related to oxidative damage due to its similarity to the *MUTYH* signature, there was no other evidence to support it. The observations about the co-occurrence of Signatures 18 and 30 and their association with dysfunction of genes involved in repair of oxidative damage in other cancers fill this gap and strongly suggest that both signatures are related to oxidative damage possibly caused by oxidative stress. While the occurrence of 8-oxo-G lesions in normal cells is estimated to be around 10^3^ per cell/per day, it can be much higher in cancer cells [9]. In the situation of sustained oxidative stress, *MUTYH*-related pathways might not be able to correct all 8-oxo-G lesions even when it is functional. Importantly, while 8-oxo-G is the most abundant lesion in normal cells, oxidative stress-induced mutagenesis occurs often at cytosines [14] explaining the simultaneous presence of both signatures.

Finally, we also observe enriched transitions between Signature 18 and the DNA mismatch repair (MMR) signature 20. This is consistent with the growing understanding that the MMR pathway is also important for the response to oxidative damage. In fact, mismatch repair deficient mice show susceptibility to oxidative stress-induced intestinal carcinogenesis [38]. A study by Colussi et al. [10] showed that baseline 8-oxoG levels were higher in DNA extracted from *MSH2* and *MLH1* deficient cell lines.

These observations indicate that the analysis of transition probabilities can be extremely valuable in shedding light on the etiology of less understood signatures.

### 3.4 Evaluation against clinical and demographic data

To show the utility of our model in the clinical setting, we evaluate the assignment of mutations to their underlying signatures using clinical and demographic data. Our analysis is based on the intuition that more accurate exposures will have higher correlation with clinical and demographic data, since multiple signature exposures have been shown to correspond to exogenous factors such as a patient’s age at diagnosis [1].

We analyze the correlation between each of the signatures and five different clinical/demographic features: age, tumor grade, and final estrogen-receptor (ER), progesterone-receptor (PR), and HER2 status (Figures 5 and S4). Importantly, we separate exposures with respect to sky and clouds. This allows us to isolate clinical features that are correlated with exposures in clouds from those that result from exposure in sky. First, we evaluate the exposures of signatures with known etiologies that match our clinical dataset. For example, Signatures 1 and 5 have been hypothesized to be active in normal cells and “clock-like” due to their correlation with the age of the patient [1]. Reassuringly, we find statistically significant association between Signature 1 and 5 exposure and age, and only find these correlations for mutations in sky. As another example, Signatures 2 and 13 display patterns of mutations linked to APOBEC proteins and are correlated with APOBEC activity, which has been linked to HER2 expression in breast cancers [41, 7, 27]. Our results also capture this relationship, with statistically significant associations between exposures to both APOBEC signatures and HER2 status in both sky and clouds. The negative correlation between Signature 3 and HER2 is consistent with the observation that while BRCA1 and BRCA2 mutation carriers display homologous recombination repair deficiency, that is the phenotype underlining Signature 3, these patients are typically HER2 negative [16].

**Fig. 5.**
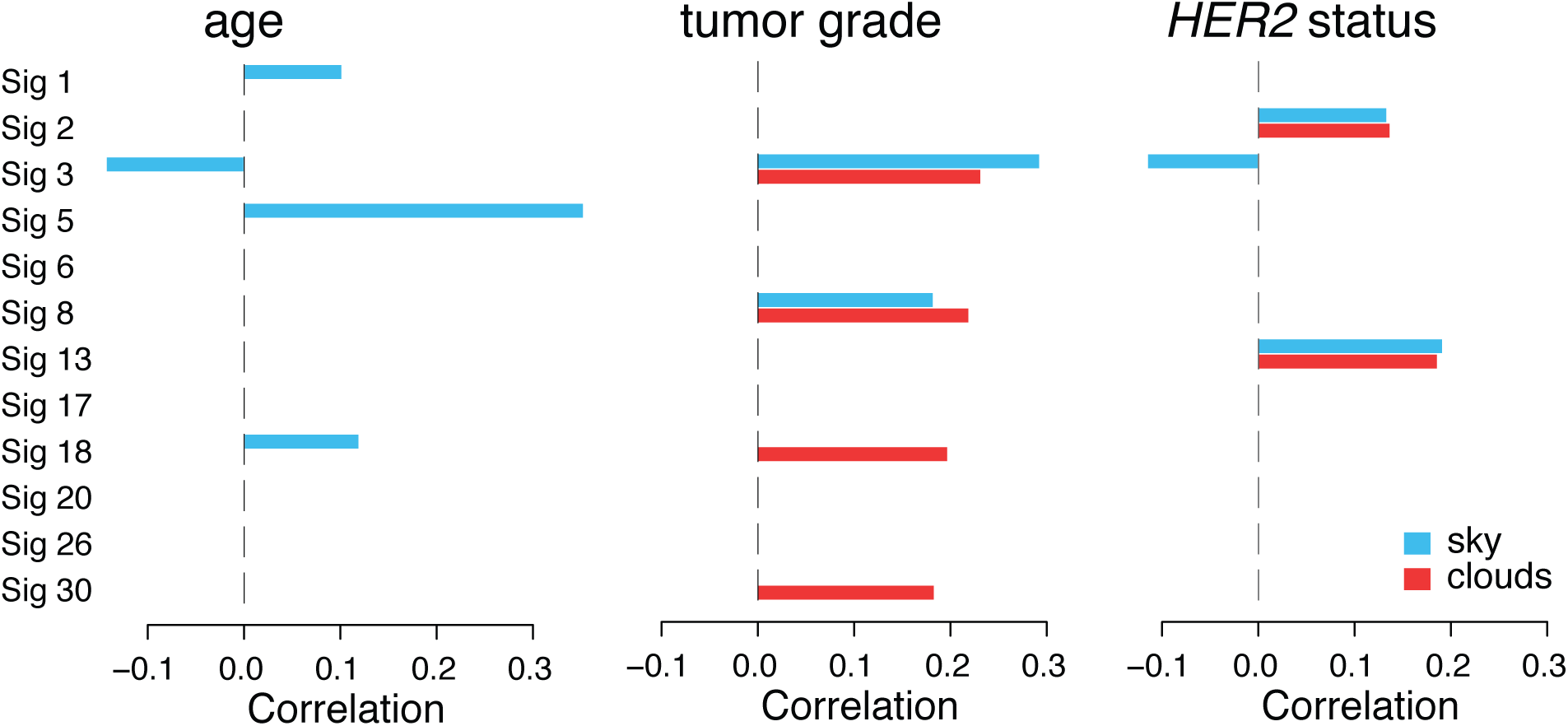
Pearson correlation coefficients between demographic or clinical features and signature exposures in sky and cloud regions. Only significant correlations with an empirical FDR *p*-value cut-off of 0.05 are shown.

We also uncover other significant associations between mutation signature exposures and age and tumor grade. Interestingly, we find that exposure to Signature 18 is significantly positively correlated with age. Coupled with our evidence that Signature 18 is involved in oxidative damage (see Section 3.3), this correlation is supported by previous observations that oxidative damage may increase with age [21]. We also find significant associations of exposure to Signature 3, 8, 18, and 30 and tumor grade. Interestingly, for Signatures 18 and 30, these associations are only statistically significant for mutations in clouds. While of unknown biological significance, these results underline the importance of the clouds in quantifying the exposure to different signatures. We report additional significant correlations in Figure S4.

Finally, we compare the overall correlation of the exposures to the 12 signatures computed with our model and NMF with the clinical and demographic data, taking overall exposures into account for both models. To this end, we compute a single correlation using canonical correlation analysis (CCA) [24]. The obtained correlation was higher for SigMa than NMF (0.674 vs. 0.665). These results provide further evidence that by using sequential information, SigMa is better able to assign mutations to signatures compared to previous models.

## 4 Discussion and conclusions

We presented the first probabilistic model of sequential dependency for mutation signatures, SigMa. We first showed that models of sequential dependency of mutation signatures have greater predictive power for held-out data than models that ignore this dependency. Next, we found that by modeling sequential dependencies previously observed among mutations [33, 31, 22, 45], we improved the estimation of mutation-to-signature assignment, and revealed new insights into the genomic factors that bias mutational process activity.

The ability to correctly determine which mutational processes generated a specific mutation is of primary importance for understanding of the emergence of tumors. For example, previous studies provided evidence that APOBEC activity is responsible for the generation of helical domain hotspot mutations in the *PIK3CA* gene in papilloma virus-driven tumors [23]. Computational tools like SigMa provide the means for finding such relationships between mutational processes and gene-level cancer drivers. A more precise assignment of mutation to signatures also allows for a more precise estimation of signature exposures and, consequently, can help to uncover relations between mutational processes and clinical and demographical phenotypes that might be difficult to infer if the signature exposure is low and signature assignment noisy. For example, our results show, for the first time, that there is a correlation between Signature 18 linked to oxidative damage and patient age. While this observation is new, it is consistent with the general understanding that an age-related increase in oxidative DNA lesions should be expected and results consistent with this explanation have been reported in few studies [21, 26]. However, the interplay of oxidative stress and cancer remains less understood [20]. The ability to correctly map oxidation related lesions might prove to be fundamental for progress in understanding the role of this process in breast cancer.

Our analysis also reinforces the idea that cloud (close-by) mutations have distinct properties from sky (isolated) mutations. While some of differences between these two mutation groups have been appreciated before (see, e.g., [45]) our analyses bring novel insights. For example, our results show that the correlation with age is a unique property of sky mutations. Interestingly, mutations that are assigned to the same signatures can have distinct properties when localized in clouds versus sky suggesting that they correspond to different subpopulations. These subpopulations, despite being assigned the same signature, might correspond to different combinations of causes. As a case in point, we found that APOBEC-associated mutations have different properties with respect to replication time depending on their assignment to sky versus clouds. As another example, while Signature 18 exposure in sky correlates with age, exposure to the same signature in clouds correlates with tumor grade.

The basic HMM model presented here can be extended and refined in various ways. In this work, we focused on modeling sequential dependency of previously validated mutation signatures from COSMIC [18]. One extension to our model is learning signatures and transitions simultaneously across multiple samples. Another possible refinement is to cast it in a Bayesian framework and add prior distributions to the model parameters. This refinement will be especially important when training the model on different cancer types where the number of samples is low.

While evaluating the predictions of SigMa using clinical and demographic data, we found a statistically significant anti-correlation of Signature 3 activity (associated with homologous recombination repair deficiency [HRD]) and patient age. We hypothesize that this is in part a consequence of germline variants predisposing to HRD (such as *BRCA1* mutations; see [40]) leading to earlier onset of breast cancer. In fact, the correlation between Signature 3 activity and age drops from −0.14 to −0.09 when removing patients Nik-Zainal, et al [34] identified as having *BRCA1* or *BRCA2* germline variants. Thus, in general, mutation signatures whose activity is anti-correlated with age may indicate that the signature’s etiology includes predisposing germline variants.

## Acknowledgements

XH, DW, YK, and TMP are supported by the Intramural Research Programs of the National Library of Medicine (NLM), National Institutes of Heath, USA. MDML gratefully acknowledges Peter Park, Jennifer Listgarten, and Nicolo Fusi for discussions regarding mutation signatures, and Michael Hoffman for discussions regarding dynamic Bayesian networks.

## Footnotes

4 https://cancer.sanger.ac.uk/cosmic/signatures

5 https://www.encodeproject.org

